# Scaffold-free cryopreservable cartilage grafts obtained from hiPSC-derived chondroprogenitor cells for airway reconstruction with growth adaptability

**DOI:** 10.1101/2025.06.05.654989

**Authors:** Shojiro Hanaki, Tomoka Takao, Yuki Fujisawa, Tomoyuki Ota, Tatsunori Osone, Shigeo Otake, Ryosuke Iwai, Koichi Deguchi, Kazuhiro Yamamoto, Hiroomi Okuyama, Takeshi Takarada

## Abstract

Pediatric tracheal reconstruction remains a major clinical challenge because of limited graft availability and the need for growth-adaptive materials. Current approaches, such as costal cartilage grafting and use of scaffold-based constructs, often suffer from complications including graft resorption, donor site morbidity, and poor integration. Here, we present scaffold-free cartilage grafts (chondro-plates) derived from expandable limb-bud mesenchymal cells generated from human leukocyte antigen-homozygous human induced pluripotent stem cells. These grafts are cryopreservable and maintain their hyaline cartilage phenotype after a brief pre-culture. In both rat and rabbit tracheal defect models, chondro-plates supported robust cartilage regeneration, epithelial reconstitution, and neovascularization. Importantly, in a pediatric-like growing rat model, chondro-plates preserved luminal patency and structural integrity, outperforming autologous costal cartilage. This study demonstrates a clinically viable, off-the-shelf strategy for tracheal reconstruction using scalable, immunocompatible, and growth-adaptive cartilage grafts.

## Introduction

Tracheal defects and stenosis caused by malignant tumors and stenotic diseases require airway reconstruction after resection of the affected tissues. Conventional airway reconstruction techniques, including end-to-end anastomosis and autologous costal cartilage transplantation, face several significant challenges. These challenges include postoperative complications, complex management, graft migration or resorption, and the invasiveness of autologous tissue harvesting. Moreover, the applicability of these techniques remains limited to specific clinical scenarios^1–3^.

Recent advancements in tissue engineering have led to the development of biodegradable polymer-based tissue-engineered tracheas (TETs) as potential alternatives for tracheal reconstruction. These TETs are typically fabricated by seeding chondrocytes onto biodegradable scaffolds engineered to replicate the native tracheal architecture and mechanics^4,5^. However, TETs face significant challenges, including granulation tissue formation on the luminal surface due to foreign body reactions, inflammation, and infection. These complications result in low survival rates of the constructs and poor engraftment of ciliated epithelial cells^4,6^. Additionally, gradual degradation of the initial cartilage, followed by its replacement with granulation tissue, remains a major limitation for the long-term success of TETs^4^.

Pediatric tracheal diseases, such as congenital tracheal stenosis, impose additional challenges due to the patients’ smaller anatomy and the necessity to accommodate their continuous physical growth^1,7^. Existing reconstructive techniques, which were primarily developed for adults, lack the flexibility and durability required for pediatric applications^8^. Thus, there is a critical need for scaffold-free regenerative medicine products that not only restore airway function but also adapt to the physical growth of pediatric patients, thereby minimizing the need for repeated surgeries^1,9^. In addition, to ensure clinical applicability in pediatric settings where surgical timing may be urgent or unpredictable, it is desirable that such grafts are pre-fabricated, cryopreservable, and usable after a short pre-culture period.

Recently, we established expandable limb-bud mesenchymal cells (ExpLBM) derived from human induced pluripotent stem cells (hiPSCs), which can be stably expanded under xeno-free culture conditions while maintaining a high chondrogenic potential^10^. These ExpLBM function as chondroprogenitor cells, which represent a stable and expandable cell population committed to the chondrogenic lineage. The viability of ExpLBM-derived hyaline cartilage-like tissues has been confirmed in small animal models, including immunodeficient mice and rats. Furthermore, we have developed a cell self-aggregation technique (CAT) that enables the fabrication of scaffold-free hyaline chondro-plates using ExpLBM^11^. In this study, we aimed to apply ExpLBM-derived chondro-plates to tracheal reconstruction. We sought to evaluate their potential to support scaffold-free airway regeneration while accommodating the continuous growth of pediatric airways, with additional assessment of their applicability as cryopreservable grafts for flexible clinical use.

## Results

### Fabrication and cryopreservation stability of ExpLBM-derived chondro-plates

Previously, we established a protocol to derive ExpLBM from hiPSCs and established a method for their expansion^10^. In this study, human leukocyte antigen (HLA)-homozygous hiPSCs (Ff-I14s04) were utilized to minimize immune rejection, which is a major challenge in regenerative medicine. The differentiation process and morphology of Ff-I14s04 at each stage are shown in Suppl. Fig. 1A. The derived ExpLBM stably expressed PRRX1 and SRY-box transcription factor 9 (SOX9), maintaining nearly 100% positivity during serial passages (Suppl. Fig. 1B). Flow cytometric analysis confirmed their high expression of CD90 and CD140B, which are surface markers associated with chondrogenic differentiation potential (Suppl. Fig. 1C).

To fabricate ExpLBM-derived chondro-plates, ExpLBM were subjected to the CAT using Step 1, Step 2, and Step 3 media (Suppl. Fig. 2)^11^. The chondro-plates were subsequently cryopreserved. A schematic overview of the experimental strategy, incorporating the use of cryopreserved chondro-plates as transplant material, is depicted in Fig. 1A. The chondro-plates were cryopreserved for 2 months and subsequently analyzed to evaluate their structural and molecular integrity after preservation. The gross appearance of a chondro-plate before and after cryopreservation is shown in Fig. 1B.

**Figure 1.**
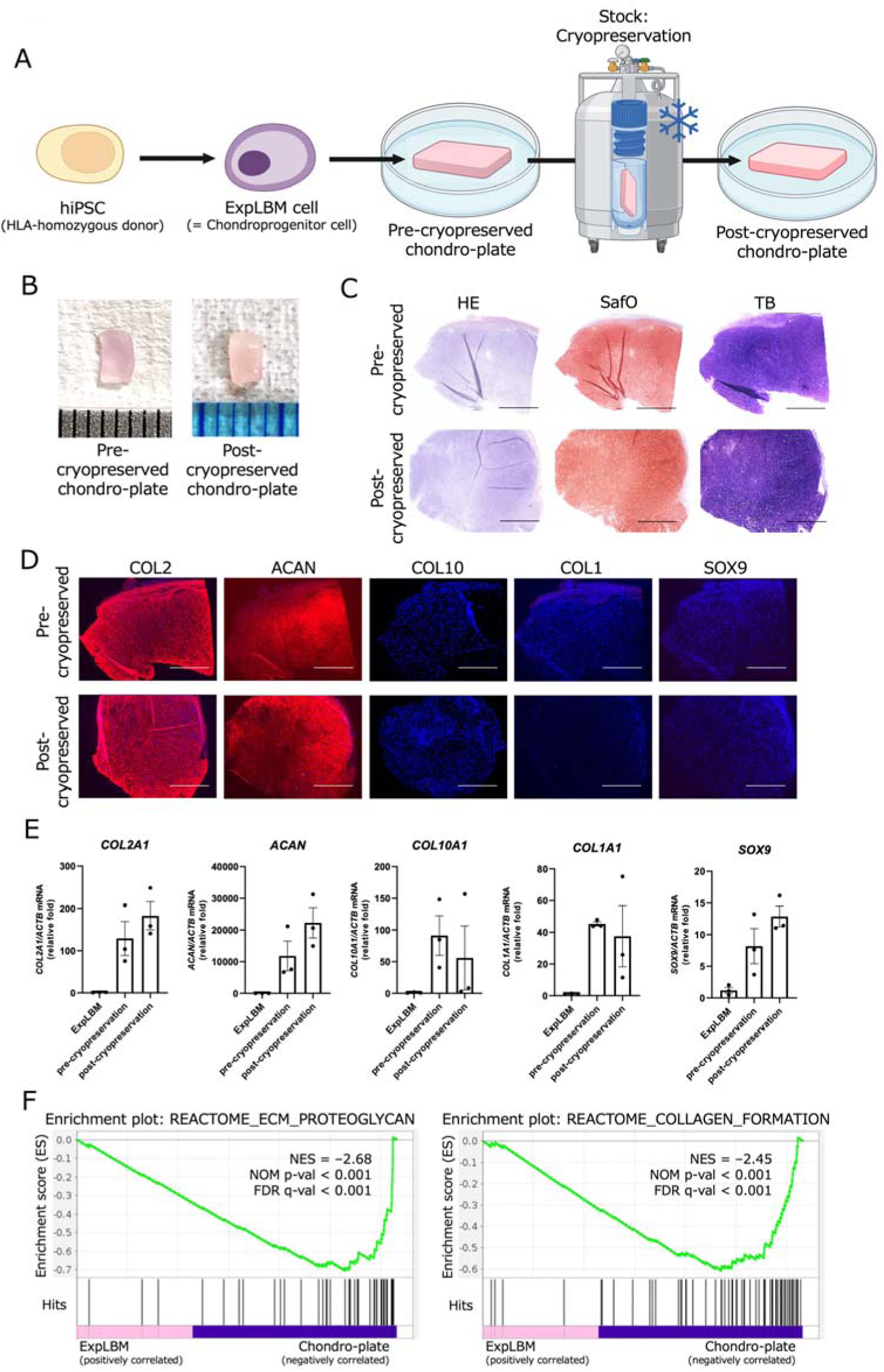
Characterization and cryopreservation of ExpLBM-derived chondro-plates. Pre- and post-cryopreserved ExpLBM-derived chondro-plates (after 2 months of cryopreservation) are compared in panels B–E. (A) Schematic diagram illustrating the differentiation process from hiPSCs to ExpLBM and fabrication of chondro-plates using the CAT. ExpLBM cells were induced from hiPSCs as previously described^10^, and chondro-plates were fabricated according to a previously reported protocol^11^. This diagram was created with BioRender.com. (B) Macroscopic appearance of chondro-plates before and after cryopreservation. (C) Histological staining of chondro-plates before and after cryopreservation. HE, SafO, and TB staining was performed to assess tissue structure and cartilage matrix production. Scale bars: 1 mm. (D) Immunohistochemical analysis of COL2, ACAN, COL10, COL1, and SOX9 expression in chondro-plates before and after cryopreservation. Scale bars: 1 mm. (E) qRT-PCR analysis of cartilage-related gene expression in pre- and post-cryopreserved ExpLBM-derived chondro-plates. Total RNA was extracted from each group, and mRNA levels of *COL2A1*, *ACAN*, *COL10A1*, *COL1A1*, and *SOX9* were quantified. All values were normalized to *ACTB* mRNA levels (n = 3, three biologically independent experiments). (F) Gene set enrichment analysis (GSEA) comparing transcriptomic profiles between ExpLBM cells and pre-cryopreserved chondro-plates. Enrichment plots are shown for extracellular matrix-related pathways, including REACTOME_ECM_PROTEOGLYCANS and REACTOME_COLLAGEN_FORMATION, which were significantly enriched in chondro-plates. In both plots, genes ranked toward the right exhibit higher expression in chondro-plates, indicating a transcriptomic shift toward a cartilage-producing phenotype.

Histological analyses demonstrated that cells were embedded in an extracellular matrix (ECM) that was uniformly stained with Safranin O (SafO) and Toluidine blue (TB) both before and after cryopreservation (Fig. 1C). Immunohistochemical staining revealed that embedded cells expressed collagen type II (COL2) and aggrecan (ACAN), whereas collagen type X (COL10), a hypertrophic cartilage marker; collagen type I (COL1), a fibrocartilage marker; and SOX9, a chondrogenic transcription factor, were not detected (Fig. 1D). No notable changes in histological or immunohistochemical profiles were observed following cryopreservation. Furthermore, gene expression analyses showed that the levels of *COL2A1*, *ACAN*, *COL10A1*, *COL1A1*, and *SOX9* expression were comparable before and after cryopreservation (Fig. 1E). Transcriptomic analysis demonstrated significant enrichment of cartilage matrix-associated pathways in pre-cryopreserved chondro-plates relative to ExpLBM cells, including ECM proteoglycans and collagen formation (Fig. 1F).

These findings demonstrate that cryopreserved ExpLBM-derived chondro-plates retain their cartilage phenotype and are suitable for use as transplantable grafts in vivo.

### Short-term engraftment and cartilage regeneration after ExpLBM-derived chondro-plate transplantation in rats

Following fabrication and cryopreservation, ExpLBM-derived chondro-plates were transplanted into tracheal defects in immunodeficient rats. A schematic overview of the transplantation procedure is shown in Fig. 2A. To assess the regenerative efficacy of these chondro-plates in vivo, defects measuring 1.5 × 2.5 mm were created in the trachea of X-linked severe combined immunodeficiency (X-SCID) rats, into which chondro-plates were implanted (Fig. 2B). All transplanted rats (n = 5) survived with stable respiratory function throughout the observation period.

**Figure 2.**
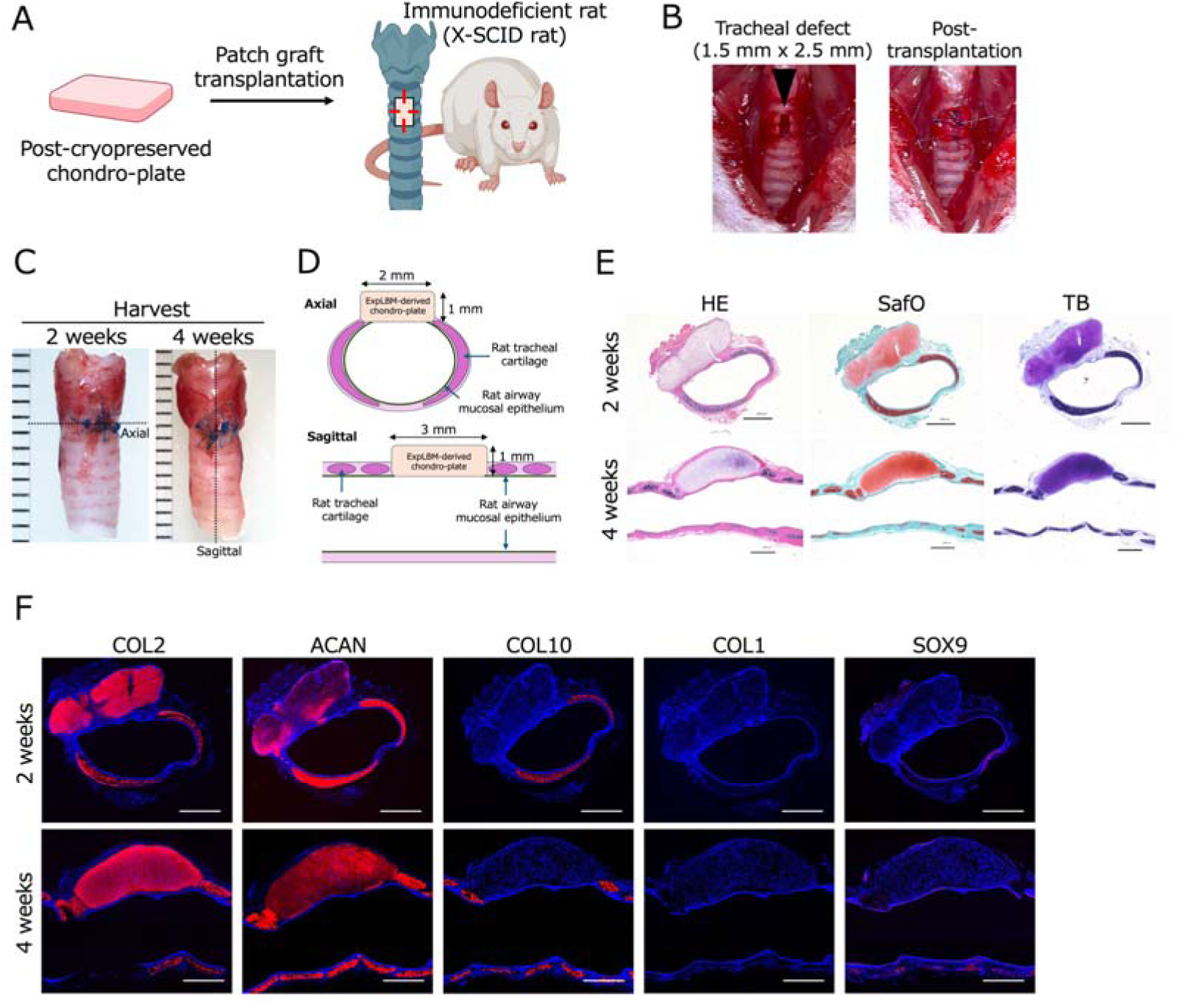
Short-term evaluation of ExpLBM-derived chondro-plate transplantation into the rat trachea. (A) Schematic representation of the transplantation procedure. A defect was surgically created in the cervical trachea of immunodeficient rats and then a post-cryopreserved ExpLBM-derived chondro-plate was implanted. This diagram was created with BioRender.com. (B) Intraoperative images of the trachea showing the surgically created defect (1.5 mm × 2.5 mm, indicated by an arrowhead) and the transplanted area after implantation of the chondro-plate. (C) Gross appearance of the transplanted trachea at 2 and 4 weeks post-transplantation. (D) Schematic illustrations indicating the positions of axial and sagittal sections used for histological evaluation. (E) Histological evaluation of transplanted chondro-plates at 2 and 4 weeks post-transplantation. HE, SafO, and TB staining was performed to assess cartilage matrix formation and integration with the host tracheal tissue. Scale bars: 1 mm. (F) Immunohistochemical analysis of the cartilage-related markers COL2, ACAN,COL10, COL1, and SOX9 at 2 and 4 weeks post-transplantation. Scale bars: 1 mm.

At 2 and 4 weeks post-transplantation, ExpLBM-derived chondro-plates were successfully engrafted into the tracheal defects (Fig. 2C). Histological sections were obtained in both the axial and sagittal planes. A schematic diagram of the sectioning planes is shown to aid interpretation of the histological images (Fig. 2D).

Histological staining revealed that transplanted chondro-plates maintained their structure and were well integrated into the surrounding tracheal tissues. Hematoxylin and eosin (HE) staining showed that cells were uniformly distributed within the grafts, while SafO and TB staining indicated that cartilage-specific ECM was consistently deposited (Fig. 2E). No evidence of graft dislocation or a severe inflammatory response was observed at either time point.

Immunohistochemical analysis demonstrated that chondro-plates exhibited strong positivity for COL2 and ACAN, confirming cartilage matrix production, whereas COL10, a hypertrophic marker, was minimally detected (Fig. 2F). COL1, a fibrocartilage-associated marker, and SOX9, a transcription factor essential for chondrogenesis, were not observed by immunostaining at either time point.

Additionally, engraftment of human-derived cells was confirmed by immunostaining for human vimentin, which was predominantly detected within the transplanted region at both 2 and 4 weeks post-transplantation (Fig. 3A, B). Regeneration of the host tracheal ciliated epithelium was evaluated by acetylated α-tubulin staining. At 2 weeks, re-epithelialization was observed, with only minimal focal discontinuities noted along the luminal surface of the graft (Fig. 3B, panel a). By 4 weeks, the epithelial lining was fully restored over the transplanted region (Fig. 3B, panel c), indicative of progressive and complete re-epithelialization. Furthermore, CD31 staining demonstrated neovascularization on the surface of transplanted chondro-plates, indicative of early vascular association with the graft (Fig. 3C).

**Figure 3.**
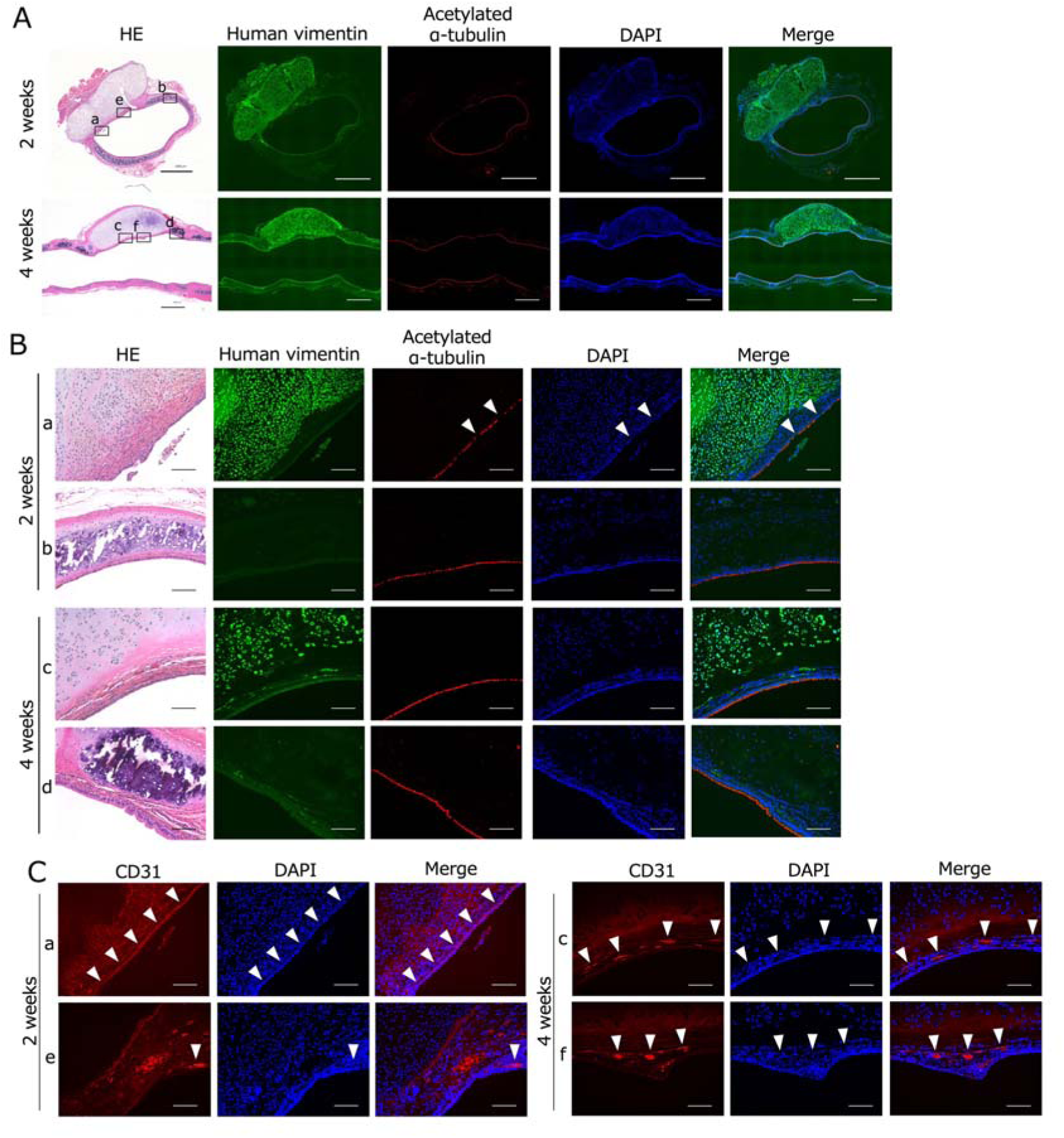
Immunohistochemical analysis of human cell engraftment and airway regeneration after ExpLBM-derived chondro-plate transplantation. (A) Immunohistochemical staining of human vimentin and acetylated α-tubulin at 2 and 4 weeks post-transplantation to evaluate engraftment of human-derived cells and reconstruction of ciliated epithelial structures. Scale bars: 1 mm. (B) Higher magnification images corresponding to (A), showing the transplanted area (a, c) and adjacent normal tracheal tissue (b, d) for comparison. In (a), partial discontinuity of acetylated α-tubulin-positive ciliated structures is observed (arrowheads). Scale bars: 100 μm. (C) Immunohistochemical staining of CD31 at 2 and 4 weeks post-transplantation. Higher magnification images of the transplanted area are shown. Panels a, c, e, and f represent different regions within the grafted tissue demonstrating CD31-positive neovascularization (arrowheads). Notably, (a) shows a linear vascular pattern. Scale bars: 100 μm.

Collectively, these findings confirm that within the first 4 weeks following transplantation, ExpLBM-derived chondro-plates successfully engrafted, formed cartilage matrix, and supported epithelial regeneration and neovascularization in the rat trachea.

### Growth-associated changes in body size and tracheal dimensions in rats

To evaluate growth-associated changes in body size and tracheal dimensions, X-SCID rats were examined at four developmental stages: 2.5, 5, 10, and 20 weeks of age. Representative images of rat morphology and excised organs (trachea, heart, and lungs) at each time point are shown in Fig. 4A. Isolated tracheal specimens were macroscopically assessed to document morphological changes over time (Fig. 4B).

**Figure 4.**
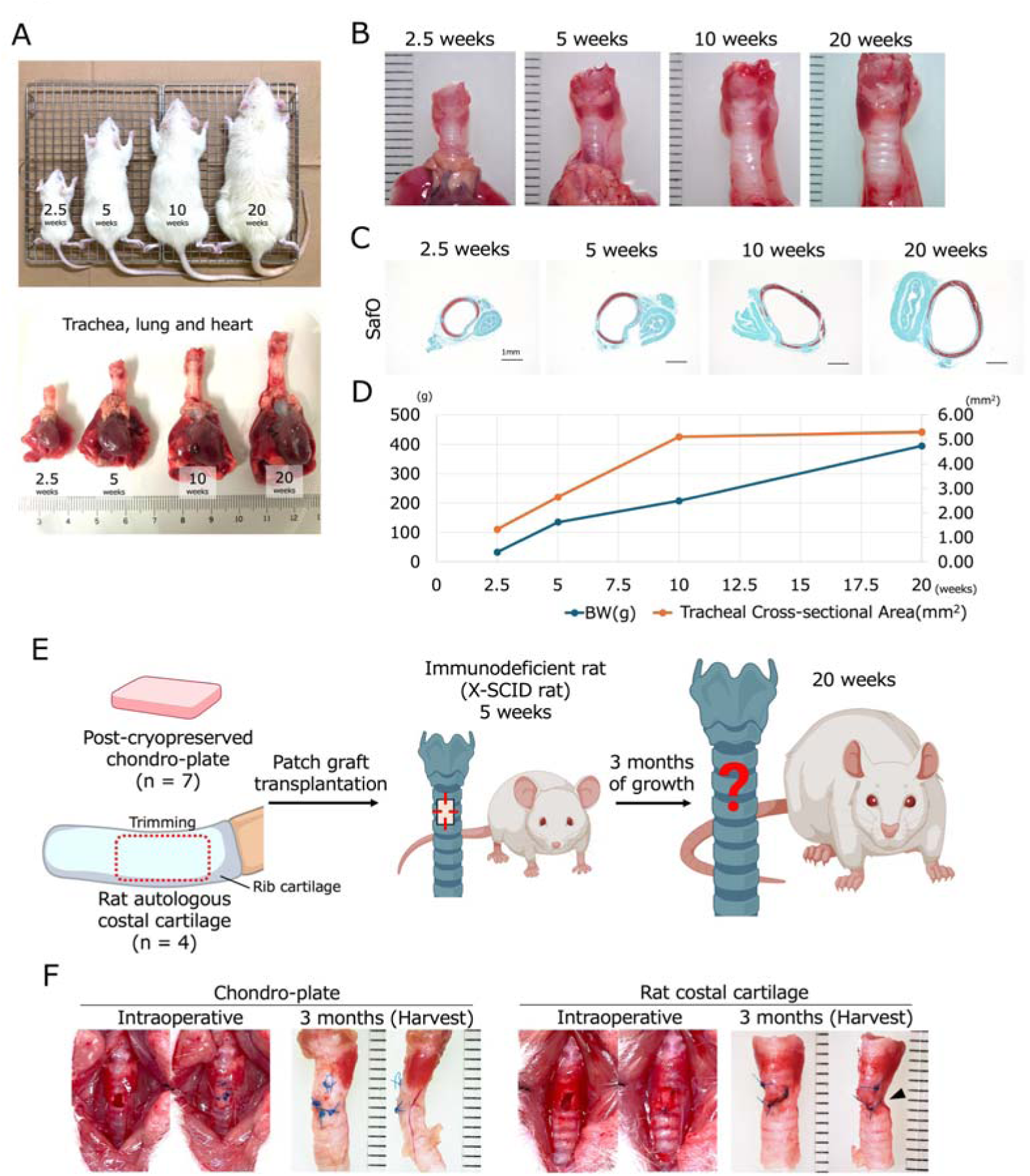
Growth-dependent changes in the trachea and evaluation of patch graft transplantation using ExpLBM-derived chondro-plates and autologous costal cartilage in rats. (A) Gross appearance of rats at different ages and macroscopic views of the excised trachea, heart, and lungs collected together after sacrifice. (B) Macroscopic appearance of the isolated trachea after separation from the heart and lungs. (C) SafO staining of transverse tracheal sections at the intended transplantation site, showing structural changes at different ages. Scale bars: 1 mm. (D) Quantitative analysis of body weight and the tracheal luminal area at 2.5, 5, 10, and 20 weeks of age. Body weight was measured using an electronic balance, and the tracheal luminal area was measured using ImageJ software based on SafO-stained transverse sections. (E) Schematic diagram illustrating the long-term experimental design to assess growth adaptability following transplantation. The red dotted box indicates the rib (costal) cartilage trimming site from which the autologous graft was harvested from the same rat. This diagram was created with BioRender.com. (F) Intraoperative views during transplantation and gross appearance of the grafted trachea at 3 months post-transplantation for both graft types. Arrowheads indicate deformed and depressed regions of the grafted area.

Histological analysis using SafO staining of transverse sections revealed that the tracheal lumen progressively enlarged as rats matured (Fig. 4C). Quantitative measurements demonstrated that the cross-sectional area of the trachea nearly doubled between 5 and 10 weeks of age (Fig. 4D). Thereafter, the increase of the tracheal cross-sectional area plateaued, indicating that major tracheal growth had occurred by 10 weeks of age.

These data informed the design of a transplantation protocol, in which rats underwent surgery at 5 weeks of age and were evaluated at 20 weeks of age, allowing assessment of graft adaptability during rapid somatic and tracheal growth.

### Long-term and growth-dependent evaluation of transplantation using ExpLBM-derived chondro-plates and autologous costal cartilage in rats

Using the growth-adapted transplantation protocol, we implanted ExpLBM-derived chondro-plates (n = 7) or autologous costal cartilage (n = 4) into 5-week-old rats and evaluated outcomes at 20 weeks of age. A schematic overview of the experimental plan and evaluation time points is shown in Fig. 4E. All transplanted rats survived until the scheduled endpoint.

Gross examination revealed that two of four rats in the autologous costal cartilage group exhibited visible deformities in the grafted trachea (Fig. 4F). By contrast, none of the rats in the ExpLBM-derived chondro-plate group exhibited macroscopic deformities or airway stenosis.

Histological evaluation of transverse sections was performed using HE, SafO, and TB staining (Fig. 5A). In axial section area measurement analysis of transplant sites, five rats in the ExpLBM-derived chondro-plate group and three rats in the autologous costal cartilage group were compared. The autologous costal cartilage group exhibited significant stenosis (p < 0.001). The degree of stenosis was classified as Myer–Cotton’s grade II or higher (Fig. 5B). By contrast, all seven rats in the ExpLBM-derived chondro-plate group maintained well-preserved tracheal architecture, with no evidence of deformation or stenosis. Consistent with these structural findings, SafO and TB staining revealed that cartilage matrix in the ExpLBM-derived chondro-plate group remained uniformly stained, indicative of a sustained proteoglycan content, whereas the autologous costal cartilage group exhibited irregular or reduced staining intensity in regions of deformation (Fig. 5A).

**Figure 5.**
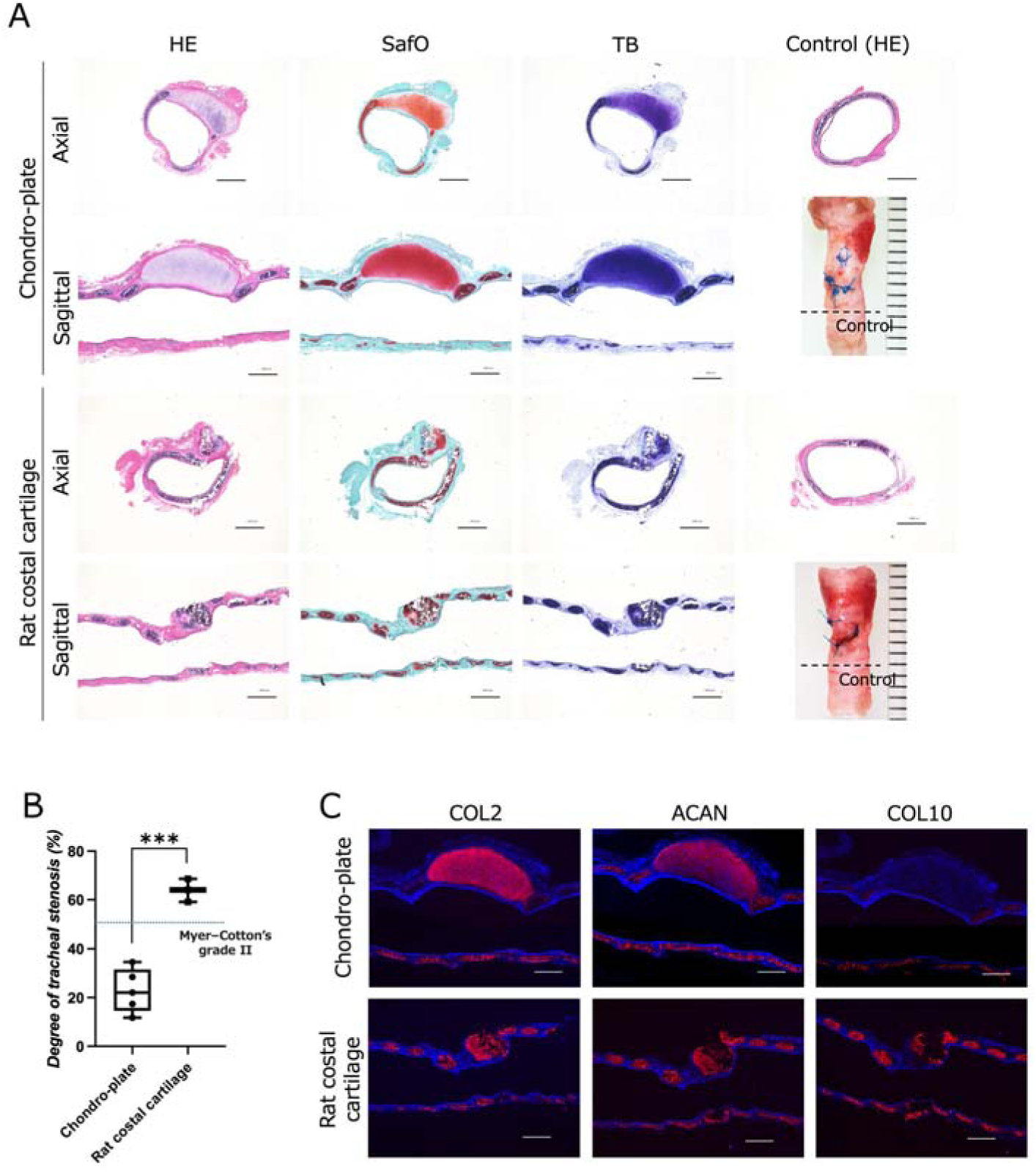
Histological and immunohistochemical comparison of ExpLBM-derived chondro-plate and autologous costal cartilage grafts at 3 months post-transplantation. (A) HE, SafO, and TB staining of axial and sagittal sections of the grafted areas at 3 months post-transplantation. Axial sections of the adjacent normal trachea were used as controls. Scale bars: 1 mm. (B) Comparison of stenosis rates between chondro-plates (n = 5) and rat costal cartilage (n = 3). The data represent the percentage of stenosis observed in each group. Statistical significance was determined using the unpaired two-tailed Student’s t-test. *** pL<L0.001. (C) Immunohistochemical staining of the cartilage-related markers COL2, ACAN, and COL10 in the grafted areas at 3 months post-transplantation. Scale bars: 1 mm.

Immunohistochemical analysis showed that both groups were positive for the cartilage-specific markers COL2 and ACAN. By contrast, COL10 expression was absent in the ExpLBM-derived chondro-plate group but was partially observed in the autologous costal cartilage group (Fig. 5C).

The persistence of human-derived cells in the ExpLBM-derived chondro-plate group was confirmed by positive staining for human vimentin (Fig. 6A). Evaluation of airway epithelial regeneration using acetylated α-tubulin staining revealed that epithelial coverage over the grafted region was limited in the autologous costal cartilage group (Fig. 6A, panel c), in contrast to the more continuous epithelial lining observed in the ExpLBM-derived chondro-plate group (Fig. 6A, panel a).

**Figure 6.**
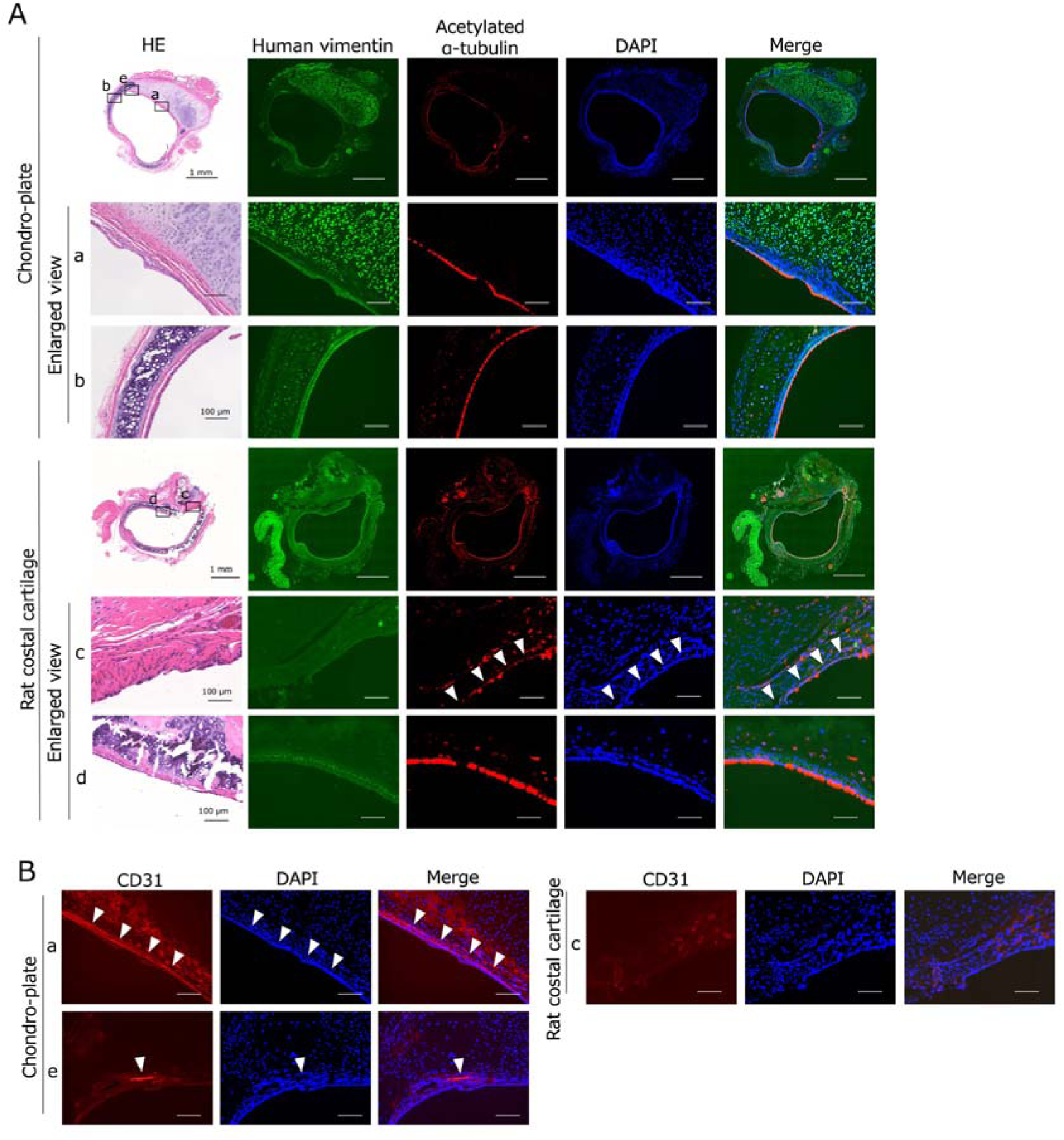
Immunohistochemical analysis of human cell engraftment, airway regeneration, and neovascularization at 3 months after transplantation of ExpLBM-derived chondro-plates and autologous costal cartilage. (A) Immunohistochemical staining of human vimentin and acetylated α-tubulin at 3 months post-transplantation to assess human cell engraftment and reconstruction of ciliated epithelial structures. Higher magnification images highlight the transplanted area (a, c) and adjacent normal tracheal tissue (b, d). In panel (c), prominent discontinuity in acetylated α-tubulin-positive ciliated structures is observed (arrowheads). Scale bars: 1 mm (low magnification) and 100 μm (high magnification panels a–d). (B) Immunohistochemical staining of CD31 at 3 months post-transplantation. Higher magnification images (a, c, e) show different regions within the transplanted area. CD31-positive neovascularization is evident in panels (a) and (e) (arrowheads), while minimal-to-no vascularization is observed in panel (c). Scale bars: 100 μm.

Furthermore, CD31 staining demonstrated neovascularization on the surface of transplanted tissues in the ExpLBM-derived chondro-plate group (Fig. 6B, panels a and e), whereas vascular structures were not clearly detected in the autologous costal cartilage group (Fig. 6B, panel c).

Collectively, these findings demonstrate that while rats in both groups survived, ExpLBM-derived chondro-plates more effectively preserved tracheal architecture, promoted epithelial regeneration, and supported neovascularization under growth-associated mechanical stress.

### Establishment of an immunosuppressed rabbit model for tracheal transplantation

To establish a rabbit model for xenogeneic tracheal transplantation, we developed a tacrolimus-based immunosuppression protocol. A schematic overview of the experimental design is shown in Fig. 7A.

**Figure 7.**
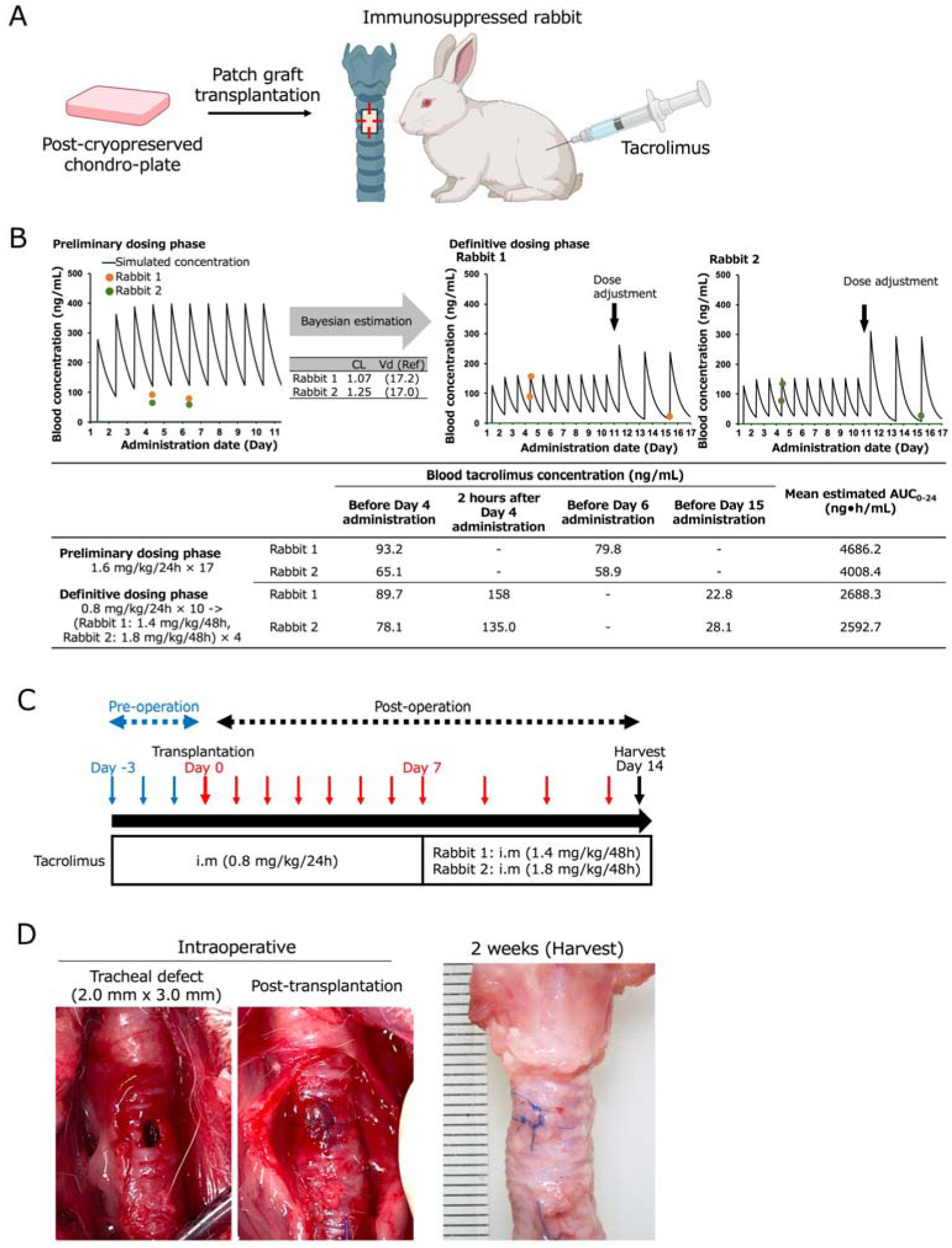
Experimental design and immunosuppressive management for ExpLBM-derived chondro-plate transplantation into the rabbit trachea. (A) Schematic diagram illustrating the overall experimental design for ExpLBM-derived chondro-plate transplantation into female JW rabbits. This diagram was created with BioRender.com. (B) PK assessment and individualized dosing strategy. In the preliminary dosing phase, tacrolimus was administered to two rabbits based on a regimen designed using reported PK parameters to evaluate individual variability. In the definitive dosing phase, tacrolimus was administered intramuscularly at individualized doses to maintain target blood concentrations. The simulated blood tacrolimus concentration curve, measured trough levels, and estimated AUC are shown. (C) The tacrolimus administration schedule following transplantation, indicating the dosing regimen maintained until sacrifice at 2 weeks post-transplantation. (D) Intraoperative view during transplantation and gross appearance of the grafted trachea at 2 weeks post-transplantation.

In the preliminary dosing phase, two female Japanese White (JW) rabbits received intramuscular tacrolimus, and blood samples were collected at defined time points to evaluate pharmacokinetic (PK) parameters. The individual systemic clearance (CL) and volume of distribution (Vd) are presented in the table embedded in Fig. 7B. Based on these results, in the definitive dosing phase, two rabbits received individualized dosing regimens to maintain target blood concentrations throughout the experimental period.

The timing of tacrolimus administration and the transplantation procedure are illustrated in Fig. 7C. Tacrolimus treatment began 3 days prior to surgery and continued until sacrifice at 2 weeks post-transplantation. Blood concentrations of tacrolimus were successfully maintained within the desired therapeutic range.

Gross examination at 2 weeks confirmed that transplantation was successful without severe complications, demonstrating the feasibility and stability of this immunosuppressed rabbit model (Fig. 7D).

### Short-term outcomes of ExpLBM-derived chondro-plate transplantation in rabbits

To assess the short-term performance of ExpLBM-derived chondro-plates in a large animal model, tracheal defects in female JW rabbits were repaired using these grafts. Gross observations at 2 weeks post-transplantation revealed no evidence of graft displacement, airway stenosis, or severe inflammatory responses.

Histological analyses were performed of both axial and sagittal sections of the grafted trachea. HE staining demonstrated that transplanted chondro-plates were well integrated with adjacent tracheal tissue and maintained their structural integrity (Fig. 8A). Consistent with previous findings, SafO and TB staining confirmed the presence of a well-developed ECM within the graft (Fig. 8A).

**Figure 8.**
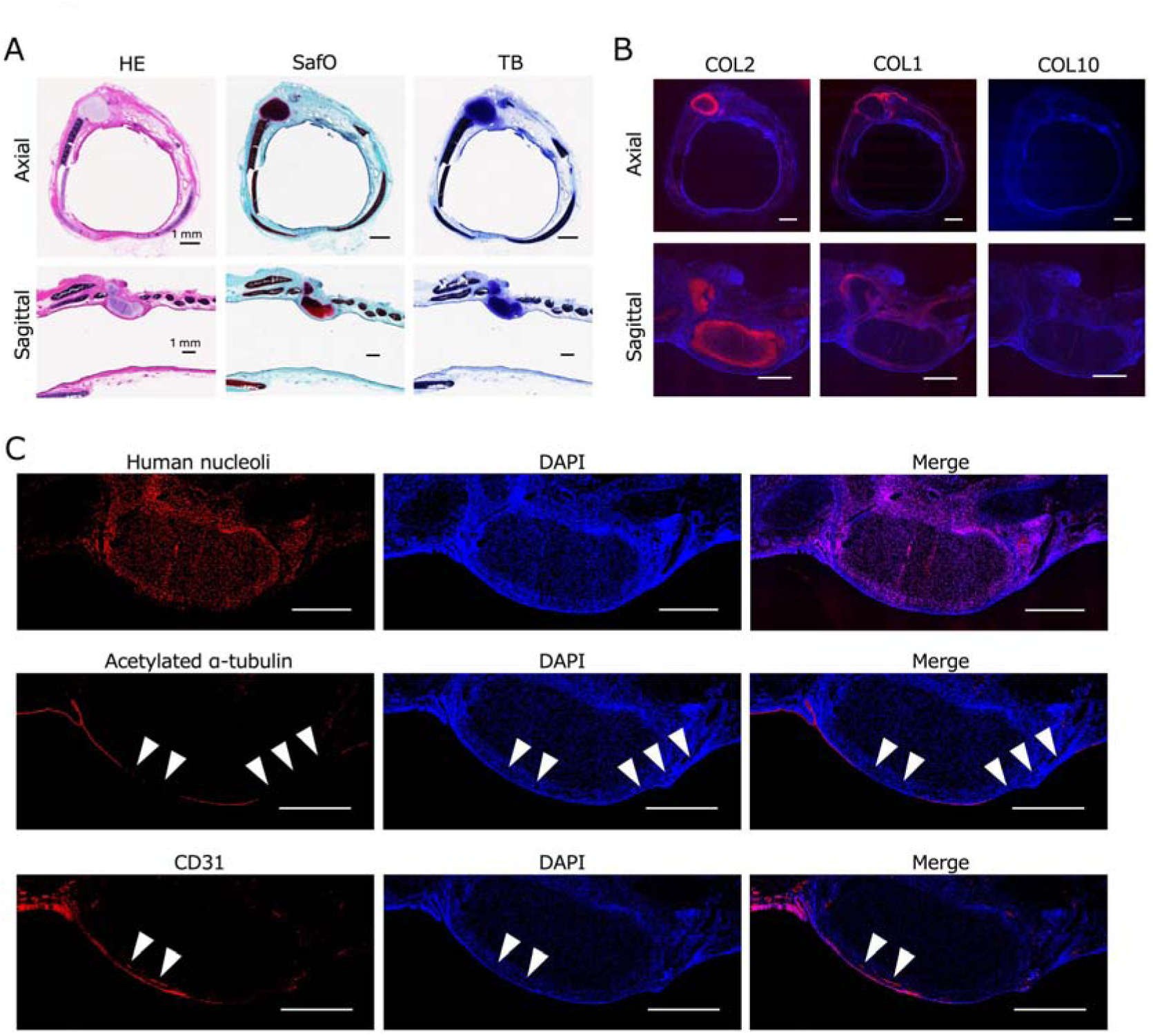
Histological and immunohistochemical evaluation following ExpLBM-derived chondro-plate transplantation into the rabbit trachea. (A) HE, SafO, and TB staining of axial and sagittal sections of the grafted region at 2 weeks post-transplantation, demonstrating overall tissue architecture and cartilage matrix formation. Scale bars: 1 mm. (B) Immunohistochemical staining of COL2, COL1, and COL10 in axial and sagittal sections to assess the cartilage phenotype and maturation status of the grafted tissue. Scale bars: 1 mm. (C) Human nucleoli-positive cells were detected within the graft, indicating that transplanted human-derived cells persisted. Acetylated α-tubulin staining revealed reconstructed ciliated epithelial structures, with minor discontinuities observed (arrowheads). CD31-positive microvessels were also identified on the surface of the graft (arrowheads), indicative of active neovascularization. Scale bars: 1 mm.

Immunohistochemical analyses revealed that COL2 was strongly expressed, whereas COL10, a marker of hypertrophic cartilage, was not detected (Fig. 8B). Collagen type I (COL1) was mainly located at the graft periphery, suggesting that fibrous tissue formed at the graft–host interface.

Staining for human nucleoli confirmed the persistence of human-derived cells within the graft. Evaluation of airway epithelial regeneration showed re-epithelialization over the grafted surface, with positive staining for acetylated α-tubulin indicating the presence of ciliated epithelial structures (Fig. 8C). Additionally, CD31 staining demonstrated neovascularization on the surface of the graft, reflecting early vascular integration (Fig. 8C).

Collectively, these findings indicate that within 2 weeks of transplantation, ExpLBM-derived chondro-plates preserved cartilage identity, promoted epithelial regeneration, and supported vascularization in rabbits.

## Discussion

This study presents two major advances in scaffold-free tracheal reconstruction. First, we demonstrate that scaffold-free cartilage grafts, fabricated as chondro-plates from HLA-homozygous hiPSCs, retain their function after cryopreservation and require only a brief pre-culture period prior to use. This property enables their application as off-the-shelf products for immediate clinical use. Second, we report the first successful evaluation of tracheal transplantation in a growth-associated model that simulates pediatric airway expansion. The chondro-plates retained anatomical integrity without deformation or stenosis, demonstrating better performance than autologous costal cartilage grafts in vivo. When transplanted into tracheal defects in immunodeficient rats, they exhibited stable engraftment, robust cartilage matrix regeneration, and both epithelial and vascular integration. Additionally, transplantation into immunosuppressed rabbits confirmed the short-term feasibility and scalability of this approach in a large animal model. Collectively, these findings highlight the clinical potential of ExpLBM-derived chondro-plates as scalable, off-the-shelf grafts for pediatric tracheal reconstruction.

Tracheal reconstruction is often necessary in patients with malignant tumors, trauma, or stenotic airway diseases. Autologous costal cartilage reconstruction remains a commonly performed procedure, but its application is limited by risks including vascular erosion, infection, limited availability, tracheomalacia, calcification, fibrosis, deformation, and loss of graft integrity^12,13^. Scaffold-free tissue engineering has emerged as a promising alternative to conventional scaffold-based methods, which often face challenges such as inflammation, foreign body reactions, and mechanical degradation over time^14,15^. In the context of tracheal reconstruction, scaffold-based TETs have demonstrated limited clinical success, primarily due to granulation tissue formation, impaired epithelialization, and compromised mechanical stability^4–6,16,17^. On the other hand, decellularized tracheal grafts offer a potential new option for tracheal regenerative medicine. One study reported that rabbits survived for up to 1 year following decellularized trachea transplantation^18^. However, if decellularization is incomplete, scaffolds can ultimately degrade and fail^14,19,20^. Additionally, issues such as granulation tissue formation and obstruction or degradation due to weakened strength during the decellularization process have also been identified^21^. To overcome these limitations, we employed a CAT to fabricate chondro-plates composed entirely of cells and their ECM, eliminating the need for artificial scaffolds^11^. This approach not only avoids scaffold-related complications but also allows the creation of mechanically stable, biologically active grafts with enhanced tissue integration. Similar to our approach, scaffold-free tracheal-like tissues were developed by creating ring-shaped cartilage using a CAT, followed by subcutaneous transplantation, and survival for up to 8 months post-transplantation was reported^22–24^. Although this method requires the invasive step of subcutaneous transplantation, it is considered to provide excellent tissue integration and strength. Previous studies attempted to use three-dimensional bioprinting or scaffold-free ring-shaped cartilage constructs for airway reconstruction; however, external stents were required or strength was insufficient in many cases^25–27^. By contrast, the chondro-plates developed in our study demonstrated sufficient durability and structural integration in vivo, supporting their applicability as a robust, scalable, and clinically translatable scaffold-free solution for airway reconstruction. Transcriptomic profiling of chondro-plates further corroborated these findings. Gene set enrichment analysis demonstrated significant enrichment of cartilage-associated pathways—including ECM proteoglycans and collagen formation—relative to ExpLBM cells, supporting the acquisition of a cartilage-like molecular identity. Traditional cartilage regeneration typically relies on autologous chondrocytes as the cell source^28–30^. Such autologous transplantation offers the advantage of high chondrogenic potential without the need to consider immune responses. However, it also presents challenges such as the limits of expansion culture and the invasiveness of autologous tissue harvesting. Furthermore, it remains challenging to obtain a sufficient number of these cells^31^. Recently, stepwise induction methods to generate cartilage from hiPSCs have been developed, and these innovative techniques can be used to produce large quantities of high-quality cartilage^32^. However, most of these methods lack intermediate cells that can be expanded or stored, limiting their use in various tissue engineering applications. ExpLBM used in this study, which are derived from hiPSCs using a defined stepwise differentiation protocol, offer several advantages as a cell source for cartilage regeneration^10,33^. Unlike mature chondrocytes, which are limited in number and exhibit a reduced proliferation capacity in vitro, ExpLBM maintain high chondrogenic potential and can be stably expanded under xeno-free conditions. They are also amenable to long-term cryopreservation without loss of phenotype or functionality, facilitating scalable production of transplantable grafts. Their reproducibility and scalability make ExpLBM a practical and clinically relevant source for generating engineered cartilage, especially when large-scale production is needed for airway reconstruction.

Use of HLA-homozygous hiPSC lines is a key strategy to mitigate immune rejection in allogeneic regenerative therapies. In this study, we employed ExpLBM derived from an HLA-homozygous hiPSC donor line (Ff-I14s04), enabling generation of chondro-plates with reduced immunogenicity. A single HLA-homozygous hiPSC line can be partially matched with a substantial portion of the population. This enables a practical route for development of off-the-shelf, immunologically compatible grafts^34,35^. Minimization of HLA mismatch not only improves graft survival and long-term function but also reduces reliance on systemic immunosuppression, which is a particularly important consideration for pediatric patients who are more susceptible to the adverse effects of immunosuppressive therapy. The successful engraftment and low inflammatory response observed in both rat and rabbit models underscore the clinical feasibility of this immunological strategy in regenerative airway applications.

A critical requirement for clinical translation of tissue-engineered grafts is cryopreservability without functional loss. In this study, ExpLBM-derived chondro-plates were cryopreserved for at least 2 months and subsequently cultured for only 7 days before implantation. The 2-month cryopreservation period was selected as a representative timeframe to evaluate the feasibility of medium-term storage under clinically relevant conditions. This duration aligns with the practical needs of off-the-shelf, pre-manufactured grafts intended for flexible surgical scheduling and potential emergency use. The grafts maintained key structural and molecular characteristics, including uniform ECM deposition and robust expression of cartilage markers such as COL2 and ACAN, even after extended cryopreservation and thawing. In addition, COL10, a hypertrophic marker, and COL1, a fibrocartilage-associated marker, were not detected by immunostaining, suggesting preservation of a stable hyaline cartilage phenotype. Notably, this minimal culture step was sufficient to restore graft readiness, underscoring the feasibility of a clinically viable workflow. Pre-fabricated chondro-plates can be prepared in advance, cryostored, and reactivated by short-term culture prior to surgery. This workflow enables a truly off-the-shelf strategy with logistical flexibility that is ideally suited for both elective and emergency pediatric airway reconstructions.

Effective tracheal reconstruction demands not only structural integrity but also restoration of key functional components, including hyaline cartilage, ciliated epithelium, and vascular networks^23,24,27,36,37^. In our study, ExpLBM-derived chondro-plates demonstrated the capacity to support all three elements. Histological and immunohistochemical analyses confirmed the robust production of cartilage matrix rich in COL2 and ACAN with minimal COL10 expression, indicative of a stable, non-hypertrophic chondrogenic phenotype. In addition, neither COL1, a fibrocartilage-associated marker, nor SOX9, a key chondrogenic transcription factor, was detected, supporting the maturity and phenotypic stability of the regenerated cartilage. The luminal surface of grafts was progressively covered with acetylated α-tubulin-positive ciliated epithelium, indicating that epithelial regeneration progressively occurred within 4 weeks of transplantation. Concurrently, CD31-positive vascular structures were observed at the graft–host interface, demonstrating early neovascularization. Together, these findings demonstrate the comprehensive regenerative capacity of ExpLBM-derived chondro-plates and underscore their suitability as functional grafts for airway reconstruction.

A key challenge in pediatric airway reconstruction is the need for grafts that can accommodate dynamic tracheal growth. In this study, we established a growth-associated rat model and demonstrated that the tracheal luminal area nearly doubled between 5 and 10 weeks of age. Under these conditions, ExpLBM-derived chondro-plates maintained their structural integrity, shape, and patency over 3 months without evidence of deformation or stenosis. By contrast, autologous costal cartilage grafts exhibited a higher frequency of luminal narrowing and structural deformities, likely due to their limited capacity to adapt to the expanding airway. The observation of Myer–Cotton’s grade II or higher stenosis in the autologous costal cartilage group indicates that the airway was partially obstructed, suggesting that timely therapeutic intervention is needed. Grade II stenosis, defined as 50–70% luminal obstruction, can significantly impair respiration and often necessitates early intervention^38–41^. These findings suggest that the biomechanical and biological properties of ExpLBM-derived chondro-plates confer superior adaptability to somatic growth, which makes them particularly suitable for pediatric applications where long-term graft compatibility with anatomical changes is essential.

Bridging the translational gap between rodent models and human-scale clinical applications remains a critical challenge. To address this, we established an immunosuppressed rabbit model to evaluate the short-term outcomes of ExpLBM-derived chondro-plate transplantation in a large airway. Using tacrolimus monotherapy, we successfully maintained stable immunosuppression, enabling engraftment and evaluation of human-derived constructs in a xenogeneic setting. At 2 weeks post-transplantation, the chondro-plates demonstrated stable integration with host tracheal tissue, preserved the cartilage matrix composition, and supported regeneration of ciliated epithelial structures. CD31-positive capillary networks were also detected at the graft surface, indicative of active neovascularization. These outcomes closely paralleled those observed in the rat model and validated the capacity of the chondro-plates to maintain their structure and function in a physiologically larger and anatomically more complex environment. This rabbit model provides a critical translational platform for refining surgical techniques, evaluating immune compatibility, and verifying the scalability and feasibility of ExpLBM-based grafts in clinically relevant airway reconstruction.

Although our findings demonstrate the feasibility and translational promise of ExpLBM-derived chondro-plates for airway reconstruction, several limitations should be addressed. First, the present study was limited to partial patch grafts, and circumferential or long-segment replacements, which are commonly required in severe pediatric cases, remain to be evaluated. Second, while mid-term outcomes were favorable in both rats and rabbits, long-term durability under respiratory biomechanics as well as potential risks, such as calcification and delayed immune responses, require further investigation. Additionally, although use of HLA-homozygous hiPSCs likely mitigates immunogenicity, the full extent of immune compatibility in an allogeneic human setting has yet to be confirmed. Future research should include orthotopic transplantation using ring- or tube-shaped constructs and refinement of surgical protocols. In parallel, long-term evaluations of mechanical integrity, immune tolerance, and functional regeneration are needed in large animal models that more closely approximate human airway physiology.

In conclusion, we demonstrate that ExpLBM-derived chondro-plates are a clinically viable, scaffold-free, cryopreservable graft for tracheal reconstruction. Leveraging the scalability and chondrogenic potential of hiPSC-derived ExpLBM, combined with the structural precision enabled by the CAT, we achieved stable engraftment, functional tissue regeneration, and adaptability to somatic growth in both small and large animal models. The use of HLA-homozygous hiPSCs further enhances the immunological compatibility of this approach, supporting the development of off-the-shelf allogeneic grafts. Taken together, these findings pave the way for clinical translation through orthotopic transplantation studies and long-term evaluation of pediatric airway reconstruction.

## Data availability

The datasets generated and/or analyzed during the current study are available from the corresponding author upon reasonable request.

## Acknowledgments

The authors thank the Center for iPSC Research and Application and Kyoto University for providing the hiPSC lines. Preparation of slides and staining were supported by the Central Research Laboratory, Okayama University Medical School. We also thank the members of the Department of Animal Resources, Advanced Science Research Center, and Okayama University for maintaining rats and rabbits.

This research was supported by AMED (grant nos. 18bm0704024h0001 and 20bm0404064h0001 to T. Takarada and JP24bm1123059 to T. Takao), the Kawano Masanori Memorial Public Interest Incorporated Foundation for Promotion of Pediatrics (grant no. 35-1 to T. Takao), the JST FOREST Program (grant no. JPMJFR225H to T. Takarada), and the Terumo Life Science Foundation (to T. Takarada).

## Author contributions

S.H. and T.Takarada conceived the study. S.H. and T.Takao performed data curation. S.H., K.Y., and T.Takarada conducted formal analysis. S.H., Y.F., T.Ota, T.Osone, S.O., and T.Takao carried out investigation. S.H., T.Takao, K.Y., and T.Takarada developed methodology. T.Takarada performed project administration and supervision. R.I. contributed resources. K.D. and H.O. provided surgical guidance and clinical insights. S.H. performed visualization. S.H. wrote the original draft. S.H., T.Osone, T.Takao, K.Y., and T.Takarada performed review and editing.

All authors reviewed and approved the final version of the manuscript for submission.

## Ethics approval and consent to participate

The Ethics Committee of the Graduate School of Medicine, Dentistry and Pharmaceutical Sciences, Okayama University approved the experimental protocols for the use of hiPSCs (project title: Molecular analysis of the process of human skeletal development using human iPS cells; approval number: K1707-013; date of approval: December 6, 2019).

All animal procedures involving rats were approved by the Animal Care and Use Committee of Okayama University (approval number: OKU-2023453).

All animal procedures involving rabbits were approved by the Animal Care and Use Committee of Okayama University (approval number: OKU-2024739).

## Materials and Methods

### Ethics statement

All animal experiments were approved by the Animal Care and Use Committee and the Institutional Review Board of Okayama University. Xenogeneic implantation studies were conducted using 20 male X-SCID rats (F344-Il2rgem1Iexas; The Institute of Medical Science, The University of Tokyo) and two female JW rabbits, which were immunosuppressed with tacrolimus.

### Cell culture

HLA-homozygous Ff-I14s04 hiPSCs were provided by CiRA Foundation^42^. The culture and differentiation protocols were based on previously published methods^10,33^. hiPSCs were maintained in StemFit AK02N medium (Ajinomoto) and were dissociated with TrypLE Select (Thermo Fisher Scientific) and 0.25 mM EDTA prior to reaching confluency. The cells were resuspended in StemFit supplemented with 10 µM Y-27632 (FUJIFILM Wako) and seeded (1×10^4^ cells in 2 mL) in a 6-cm culture dish coated with 8 µL of iMatrix-511 silk (human laminin-511 E8 fragment; Nippi). The medium was changed to StemFit without Y-27632 the following day and was subsequently refreshed every 2 days until the cells reached an appropriate density for passaging.

For stepwise differentiation into limb-bud mesenchymal (LBM) cells, hiPSCs (3×10^4^) were seeded in 1 mL of StemFit supplemented with 10 µM Y-27632 and 4 µL iMatrix-511 silk in a 3.5-cm dish. After 24 h, the medium was replaced with Y-27632-free StemFit. Differentiation into mid-primitive streak, lateral plate mesoderm, and LBM stages proceeded as previously described^10^.

To passage ExpLBM, LBM cells were dissociated using accutase (Thermo Fisher Scientific) and reseeded at a density of 2–4×10^5^ cells per 6-cm dish in ExpLBM medium. This consisted of CDM2 basal medium supplemented with 3 µM CHIR99021, 1 µM A-83-01, 20 ng/mL fibroblast growth factor 2 (FGF2), 20 ng/mL epidermal growth factor, 10 µM Y-27632, and 1% penicillin-streptomycin (P/S; Gibco, 15140-122). Dishes were pre-coated with 4 µg/mL human plasma fibronectin (Merck). The medium was replaced every 2 days, and cells were passaged before reaching confluency as described above.

### Immunocytochemistry

Cells cultured on dishes were fixed with 4% paraformaldehyde for 30 min at room temperature and then incubated in blocking solution consisting of phosphate-buffered saline (PBS) containing 3% normal goat serum and 0.1% Triton X-100 for 1 h at room temperature. Primary and secondary antibodies were diluted 1:200 and 1:500, respectively, in blocking solution and applied to cells for 1 h at room temperature. After antibody incubation, nuclei were counterstained with 0.1 µg/mL 4′,6-diamidino-2-phenylindole (DAPI; Thermo Fisher Scientific). Fluorescence images were acquired using a BZ-X710 fluorescence microscope (Keyence). The primary antibodies used were anti-PRRX1 (Sigma-Aldrich, ZRB2165) and anti-SOX9 (Sigma-Aldrich, AB5535).

### Flow cytometry

Flow cytometric analysis was performed as described previously^10,33^. ExpLBM were dissociated using accutase (Thermo Fisher Scientific), and 1×10^5^ cells were resuspended in 100 µL of PBS supplemented with 2% fetal bovine serum (FBS; Sigma-Aldrich, 173012). Cells were incubated on ice for 1 h with FITC-conjugated anti-CD90 and BB700-conjugated anti-CD140B antibodies (both diluted 1:200). The CD90^high^CD140B^high^ population was analyzed using a CytoFLEX S flow cytometer (Beckman Coulter).

### ExpLBM-derived chondro-plate fabrication

Chondrogenic differentiation under adhesive culture conditions was induced using the previously established CAT as described in our earlier report^11^. To prepare the culture groove, a 10-mm diameter circle was cut from a 1-mm-thick silicone sheet using a scalpel to create a silicone frame. Silicone pillars (3 mm in diameter) were fabricated from 3-mm-thick silicone sheets. The silicone devices were cleaned using an ultrasonic cleaner, sterilized by autoclaving, and positioned on PrimeSurface® culture dishes (Sumitomo Bakelite Co.) before polymer coating. The poly(N,N-dimethylaminoethyl methacrylate)-co-poly(methacrylic acid) polymer used in the CAT was synthesized as previously described^22^. The polymer solution was diluted to 12.5 µg/mL with UltraPure™ DNase/RNase-free distilled water (Thermo Fisher Scientific) and applied to the culture groove surrounded by the silicone frame and pillar. After incubation at 25 °C for 5 min, the solution was diluted three times by aspirating half the volume and adding the same volume of PBS. The remaining solution was aspirated, and the dish was washed twice with PBS.

The cartilage induction protocol was based on the method of Takao et al.^33^. ExpLBM derived from hiPSCs were dissociated with accutase and seeded at a density of 2 × 10^6^ cells/cm^2^ in 3 mL of Step 1 medium [high-glucose DMEM (FUJIFILM, 044-29765) supplemented with 3 µM CHIR99021, 10 ng/mL FGF2, 50 µg/mL ascorbic acid, 1× insulin-transferrin-selenium (ITS), 1% P/S, and 10% FBS]. After 1 h of culture and confirmation of cell attachment, an additional 3 mL of Step 1 medium was added. After 3 days, the medium was replaced with Step 2 medium [DMEM supplemented with 10 ng/mL FGF2, 50 µg/mL ascorbic acid, 30 ng/mL bone morphogenetic protein 4 (BMP4), 10 ng/mL transforming growth factor (TGF)-β1, 10 ng/mL growth-differentiation factor 5 (GDF5), 1× ITS, 1% P/S, and 1% FBS], and the cells were cultured for another 3 days. Thereafter, the aggregates were cultured in Step 3 medium [DMEM containing 50 µg/mL ascorbic acid, 30 ng/mL BMP4, 10 ng/mL TGF-β1, 10 ng/mL GDF5, 1× ITS, 1% P/S, and 10% FBS] for 42 days. The medium was changed every 3 days.

The resulting chondro-plates were trimmed to the desired size using a scalpel and cryopreserved in liquid nitrogen using Stem Cell Banker (ZENOGEN PHARMA, 11924) as the cryoprotectant. Prior to transplantation, cryopreserved chondro-plates were thawed and cultured in Step 3 medium for 7 days. Cryopreserved chondro-plates were used for all transplantation experiments. The duration of cryopreservation varied depending on the experimental schedule and was a maximum of 2 months. All samples were subjected to the same post-thaw pre-culture process before use.

### Real-time quantitative reverse transcription polymerase chain reaction (qRT-PCR)

Total RNA was extracted using an RNeasy Mini Kit (QIAGEN), and complementary DNA was synthesized using ReverTra Ace qPCR RT Master Mix with gDNA Remover (TOYOBO) according to the manufacturer’s instructions. qRT-PCR was performed using gene-specific primers on an AriaMX Real-Time PCR System (Agilent Technologies) under the following thermal cycling conditions: initial denaturation at 95 °C for 30 sec, followed by annealing at 62 °C for 30 sec and extension at 72 °C for 30 sec. Relative gene expression levels were quantified using the ΔΔCt method, with *ACTB* serving as the internal control.

The primer sequences were as follows: *COL2A1*, forward (F) 5′-CCTGAGTGGAAGAGTGGAGACT-3′ and reverse (R) 5′-TCCTTGCTCTTGCTGCTCCA-3′; *ACAN*, F 5′-GGCACAGCCACCACCTACAA-3′ and R 5′-AGCGACAAGAAGAGGACACCG-3′; *SOX9*, F 5′-AAGCTCTGGAGACTTCTGAACGA-3′ and R 5′-CGCCTTGAAGATGGCGTTGG-3′; *COL1A1*, F 5′-CCACTGCAAGAACAGCGTGG-3′ and R 5′-GTGTGACTCGTGCAGCCATC-3′; *COL10A1*, F 5′-CCCAGCACGCAGAATCCATC-3′ and R 5′-AGTGGGCCTTTTATGCCTGT-3′; and *ACTB*, F 5′-AGAAAATCTGGCACCACACC-3′ and R 5′-AGAGGCGTACAGGGATAGCA-3′.

### RNA sequencing and gene set enrichment analysis

Total RNA was extracted using an RNeasy Kit (Qiagen), and sequencing libraries were prepared using the KAPA RNA HyperPrep Kit with RiboErase (HMR) (Kapa Biosystems, USA) and the SeqCap Adapter Kit (Set A or Set B; Roche, USA), following the manufacturers’ instructions. The libraries were submitted to Azenta (Suzhou, China) and sequenced using a HiSeq 2500 platform (Illumina, USA). Raw sequence reads were extracted in FASTQ format using the CASAVA 1.8.4 pipeline.

Adapter trimming and quality filtering were performed using Fastp (v0.23.4) by removing reads shorter than 60 bases and trimming leading and trailing bases with a Phred quality score below 30. Pseudoalignment to the GRCh38 human reference genome was conducted using Kallisto (v0.48.0). Normalization was performed using the GeTMM method implemented in edgeR (v4.0.3) to account for inter-sample variation. Differentially expressed genes were identified using NOISeq (v2.46.0), applying thresholds of probability > 0.8 and |logL fold change| > 1.

All normalized gene expression data were submitted to the GSEA command line tool (v4.3.2; Broad Institute), and gene set enrichment analysis (GSEA) was performed. The Reactome gene set collection (c2.cp.reactome.v2022.1) from the Molecular Signatures Database (MSigDB) was used.

RNA-seq data for ExpLBM cells were used from our previously published dataset (NCBI Gene Expression Omnibus, GEO accession number GSE165620). Data for pre-cryopreserved chondro-plates generated in this study have been deposited under GEO accession number GSE299047.

### Transplantation of chondro-plates and autologous costal cartilage into the rat trachea

Transplantation experiments were conducted using male X-SCID rats (F344-Il2rgem1Iexas; The Institute of Medical Science, The University of Tokyo). All procedures involving rats were approved by the Animal Care and Use Committee of Okayama University (approval number: OKU-2023453). Rats were anesthetized with isoflurane delivered via an inhalation anesthesia system (induction: 4–5%; maintenance: 2–3%). A midline cervical incision was made to expose the trachea from the cricoid cartilage to the suprasternal notch. A tracheal defect measuring 1.5Lmm (width) × 2.5Lmm (length) was created caudal to the second tracheal ring. A graft, either an ExpLBM-derived chondro-plate or autologous costal cartilage, was implanted at the defect site and secured with four interrupted 7-0 polypropylene sutures (Prolene; Ethicon Inc., Johnson & Johnson).

Two experimental protocols were employed. In the first protocol, five rats aged 15–20 weeks underwent chondro-plate transplantation. Two of these rats were euthanized at 2 weeks and three were euthanized at 4 weeks post-transplantation by CO_2_ inhalation for early engraftment assessment. In the second protocol, 11 5-week-old rats underwent transplantation. Seven of these rats received ExpLBM-derived chondro-plates and four received autologous costal cartilage grafts. All animals in this cohort were euthanized at 3 months postoperatively to evaluate long-term outcomes and adaptability of the grafts to somatic and tracheal growth.

The surgical procedure in the second protocol followed the same basic technique as that in the first, although the graft material differed between the groups. For the control group, autologous costal cartilage was trimmed from the sixth to seventh rib cartilage of the same rat and shaped to match the chondro-plate dimensions. To ensure procedural consistency, a comparable amount of costal cartilage was resected in the chondro-plate group to equalize surgical invasiveness.

Following euthanasia, tracheal segments including the graft site and adjacent tissues were excised, fixed in 70% neutral-buffered formalin (FUJIFILM Wako) for 48 h, and embedded in paraffin at the Central Research Laboratory of Okayama University Medical School. Histological and immunohistochemical analyses were conducted to assess graft integration and cartilage tissue regeneration.

### Evaluation of tracheal growth in rats

To evaluate growth-associated changes in body size and tracheal dimensions, male X-SCID rats were assessed at 2.5, 5, 10, and 20 weeks of age. At each time point, one rat was euthanized for analysis. Body weight was measured using an electronic balance. To evaluate the tracheal cross-sectional area, the cervical trachea was excised post-mortem, embedded in paraffin, and sectioned transversely. The sections were stained with SafO, and digital images were captured. The luminal area of the trachea was measured using ImageJ software (National Institutes of Health). Temporal changes in body weight and tracheal area were analyzed to assess growth dynamics.

### Immunosuppressant administration in rabbits

Immunosuppression was achieved using tacrolimus monotherapy (Astellas Pharma) in female JW rabbits. The study consisted of two phases: a preliminary dosing phase, which aimed to explore the optimal dose, and a definitive dosing phase, which was designed for surgical application.

In the preliminary dosing phase, two rabbits were enrolled to evaluate the PK parameters of tacrolimus. The initial dose of tacrolimus was set at 1.6 mg/kg/24 h. This dose was calculated based on previously reported PK parameters for JW rabbits, specifically systemic CL (0.312 L/h/kg) and Vd (6.27 L/kg)^43^. Blood samples were collected at two time points: before administration on Day 4 and before administration on Day 6. Following administration in the preliminary phase, blood concentrations of tacrolimus were used to estimate individual CL and Vd values for each rabbit by Bayesian estimation. These individualized PK parameters served as the basis for determining the initial dose in the definitive dosing phase. The AUC_0–24_ at steady state was estimated to assess drug exposure. The target AUC was determined at ≥2000 h·ng/mL based on prior studies demonstrating effective immunosuppression^44^.

In the definitive dosing phase, tacrolimus dosing was subsequently adjusted according to individual CL and Vd estimated in the preliminary dosing phase to maintain blood concentrations within the desired therapeutic range. Tacrolimus was administered intramuscularly following these individualized regimens. On Day 4 of the definitive phase, tacrolimus trough concentrations and concentrations at 2 h post-dosing were measured to evaluate whether the target AUC was achieved. The dose was adjusted accordingly to attain the desired exposure in order to ensure consistent and effective immunosuppression. Whole-blood tacrolimus concentrations were measured by SRL Inc. using an electrochemiluminescence immunoassay. The quantifiable range was 0.5–40 ng/mL. If values exceeded this range, samples were diluted and re-analyzed to obtain numerical results, in accordance with the company’s standard procedure.

### Transplantation of a chondro-plate into the rabbit trachea

Female JW rabbits were used for transplantation experiments. All animal procedures involving rabbits were approved by the Animal Care and Use Committee of Okayama University (approval number: OKU-2024739). Rabbits were anesthetized using isoflurane delivered via an inhalation anesthesia system, with an induction concentration of 4–5% and a maintenance concentration of 2–3%. A midline cervical incision was made to expose the trachea from the cricoid cartilage to the suprasternal notch. A tracheal defect measuring 2.0 mm (width) × 3.0 mm (length) was created caudal to the second tracheal ring. An ExpLBM-derived chondro-plate was implanted at the defect site and secured with four interrupted sutures using 7-0 polypropylene (Prolene; Ethicon Inc., Johnson & Johnson).

Tacrolimus-based immunosuppression was initiated 3 days prior to transplantation and maintained throughout the study. Tacrolimus was administered intramuscularly based on individualized dosing regimens determined by prior PK profiling. At 2 weeks post-transplantation, rabbits were euthanized, and tracheal segments, including the graft sites and adjacent tissues, were harvested. The specimens were fixed in 70% neutral-buffered formalin (FUJIFILM Wako) for 48 h and embedded in paraffin at the Central Research Laboratory of Okayama University Medical School. Histological and immunohistochemical analyses were performed to assess graft integration and cartilage regeneration.

### Evaluation of graft integration and cartilage regeneration in rats

Graft integration and cartilage regeneration were assessed at two time points following ExpLBM-derived chondro-plate transplantation into the rat trachea: an early stage (2–4 weeks) and a long-term stage (3 months).

For early-stage evaluation, rats were euthanized at 2 and 4 weeks post-transplantation. Only animals that received ExpLBM-derived chondro-plates were included in this analysis. Paraffin-embedded tissue sections were stained with HE, SafO, and TB to assess general tissue morphology and cartilage matrix deposition. Immunohistochemistry was performed using antibodies against COL2, ACAN, COL10, human vimentin, acetylated α-tubulin, and CD31 to evaluate cartilage-specific matrix composition, engraftment of human cells, epithelial regeneration, and neovascularization.

For long-term evaluation, rats were sacrificed at 3 months post-transplantation. Chondro-plate grafts were compared with autologous costal cartilage grafts. During the 3-month period, the body weight and tracheal dimensions of recipient rats approximately doubled. To assess whether grafts accommodated this growth, gross morphological evaluation was performed to identify deformation or luminal stenosis. Paraffin-embedded sections were prepared and stained with HE, SafO, and TB. Immunohistochemical analysis of COL2, ACAN, COL10, human vimentin, acetylated α-tubulin, and CD31 was also conducted. All histological evaluations were performed using a BZ-X710 fluorescence microscope (Keyence).

### Evaluation of graft integration in rabbits

At 2 weeks post-transplantation, rabbits were euthanized and tracheal segments containing the graft sites were harvested. Paraffin-embedded tissue sections were prepared and stained with HE, SafO, and TB to evaluate general tissue morphology and cartilage matrix production. Immunohistochemical analysis was performed using antibodies against COL2, COL1, COL10, human nucleoli, acetylated α-tubulin, and CD31 to assess cartilage matrix composition, human cell engraftment, epithelial regeneration, and neovascularization. All fluorescence images were acquired using a BZ-X710 fluorescence microscope (Keyence).

### Immunohistochemistry

Tissue samples were fixed in 10% neutral-buffered formalin (FUJIFILM Wako) or 4% paraformaldehyde and subsequently embedded in paraffin. Paraffin blocks were sectioned into 4-μm-thick slices. Sections were deparaffinized and subjected to antigen retrieval by heating in 10 mM citrate buffer (pH 6.0). After retrieval, sections were treated with 1 μg/mL proteinase K and then incubated with blocking solution (1× PBS containing 3% normal goat serum and 0.1% Triton X-100) for 1 h at room temperature. The following primary antibodies were used: anti-COL2 (Thermo Fisher Scientific, MA1-37493), anti-ACAN (Proteintech, 13889-1-AP), anti-COL10 (Cosmo Bio, LSL-LB-0092), anti-COL1 (Southern Biotech, 1441-01), anti-SOX9 (Sigma-Aldrich, AB5535), anti-human vimentin (Progen Biotechnik, 10515), anti-human nucleoli (Abcam, ab190710), anti-CD31 (OriGene, AM10064SU-N), and anti-acetylated α-tubulin (Sigma-Aldrich, T7451). All antibodies were diluted 1:200, except anti-acetylated α-tubulin, which was diluted 1:1000, using the same blocking solution. Sections were incubated with primary antibodies overnight at 4 °C and then with appropriate secondary antibodies (diluted 1:400) for 1 h at room temperature. Nuclei were counterstained with DAPI, and sections were mounted using Fluoromount-G (Southern Biotech). Fluorescence images were acquired using a BZ-X710 microscope (Keyence).

### Measurement of the tracheal stenosis percentage

Digital images of tracheal sections were captured, and the luminal area of the trachea was measured using ImageJ software (National Institutes of Health). The degree of stenosis was calculated as a percentage using the following formula: degree of stenosis (%) = (1 – minimum lumen area / nearest normal area) × 100%. The minimum lumen area was defined as the smallest cross-sectional area within the grafted segment, and the nearest normal area was taken from the adjacent native tracheal segment.

### Statistical analysis

All statistical analyses were performed using Prism 8 software (GraphPad Software). Data from five and three biologically independent experiments for the ExpLBM-derived chondro-plate and autologous costal cartilage groups, respectively, are presented as mean ± standard error of the mean. Statistical significance was assessed using the unpaired two-tailed Student’s t-test. A p-value < 0.05 was considered statistically significant.

---

**Suppl. Fig. 1.**
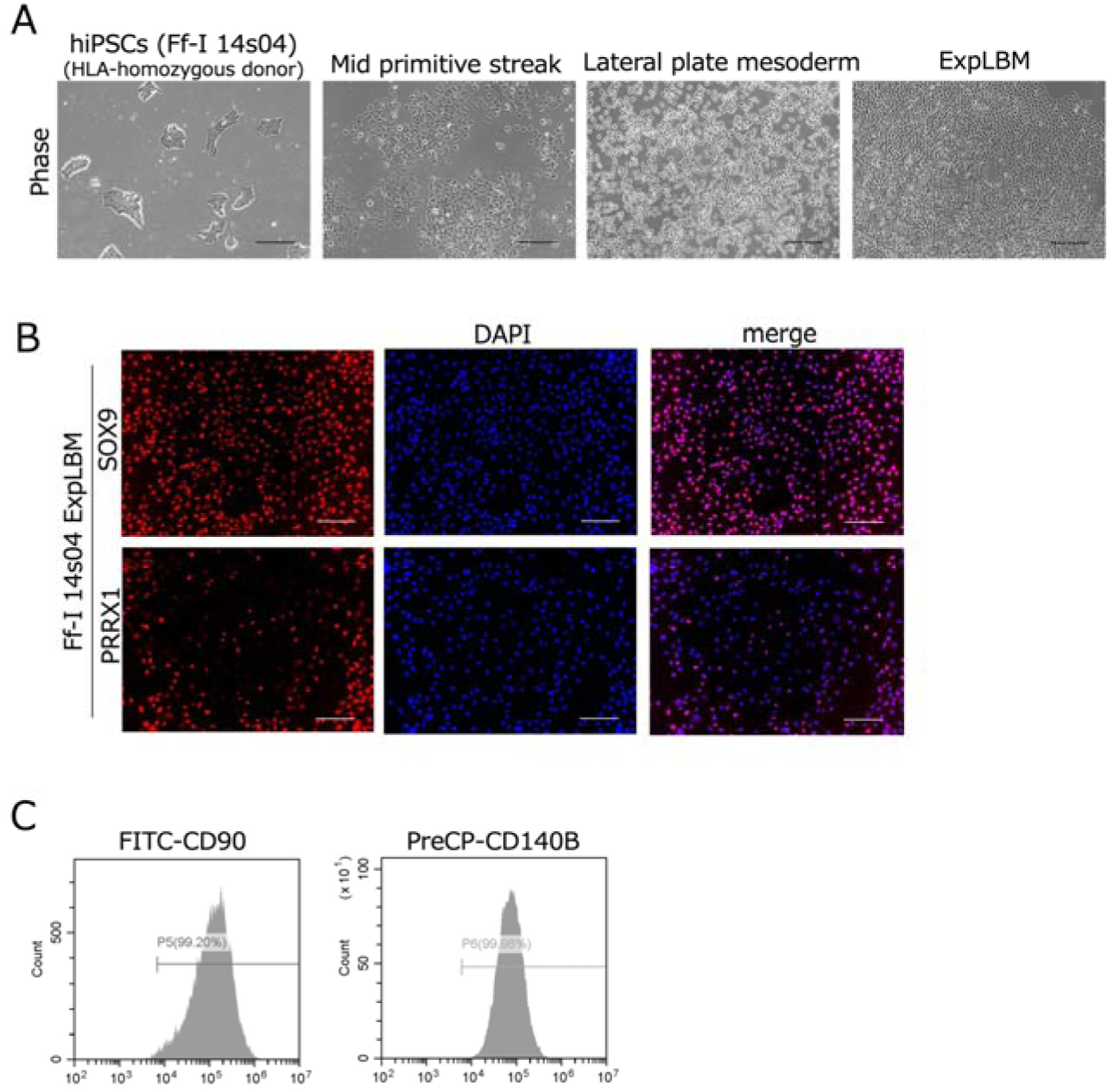
Induction and characteristics of ExpLBM. (A) Phase-contrast images showing cell morphology at each stage of differentiation from hiPSCs (Ff-I14s04) through mid-primitive streak and lateral plate mesoderm to ExpLBM. Scale bars: 200 µm. (B) Immunofluorescence staining of PRRX1 and SOX9 in Ff-I14s04-derived ExpLBM, with DAPI counterstaining. Scale bars: 200 µm. (C) Representative flow cytometric histograms of ExpLBM showing high expression of CD90 and CD140B.

**Suppl. Fig. 2.**
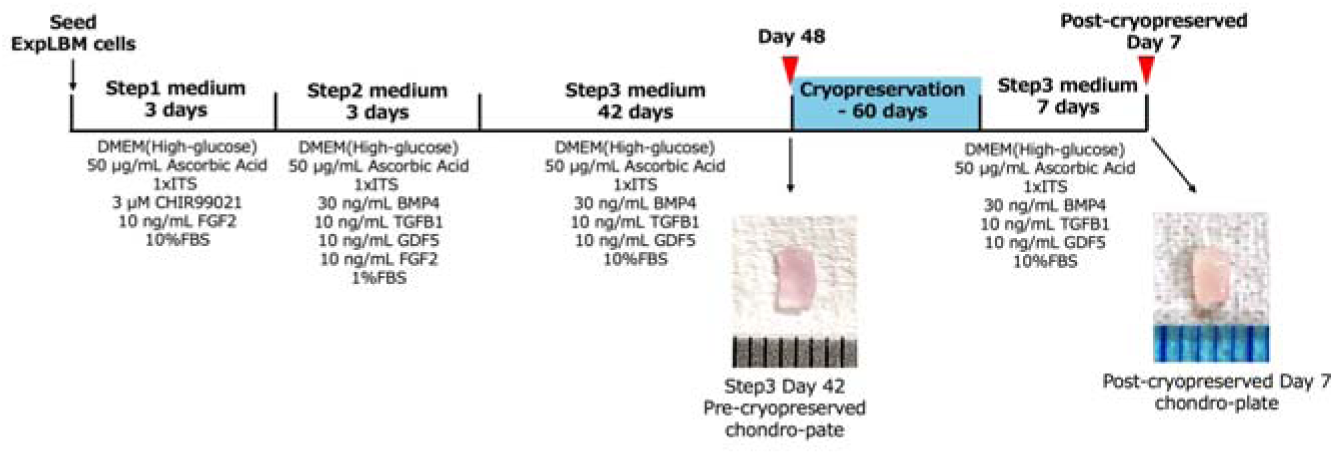
Schematic diagram of the culture protocol for induction of chondro-plates from ExpLBM using the CAT. ExpLBM were seeded and sequentially cultured in Step 1, Step 2, and Step 3 media for 48 days to generate scaffold-free chondro-plates. On Day 48, the chondro-plates were cryopreserved for 60 days. After thawing, the chondro-plates were cultured in Step 3 medium for an additional 7 days before being used for transplantation.

