## Supplemental Fig1,2 for "Scaffold-free cryopreservable cartilage grafts obtained from hiPSC-derived chondroprogenitor cells for airway reconstruction with growth adaptability"

Suppl. Fig1

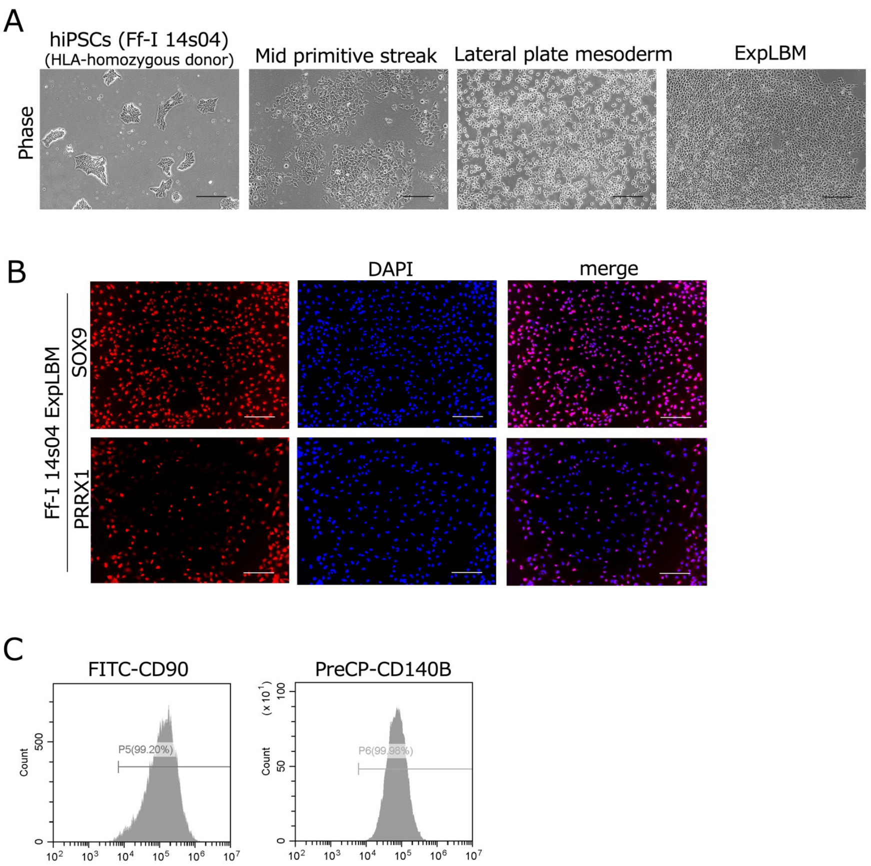


**Suppl. Fig. 1. Induction and characteristics of ExpLBM.**

(A) Phase-contrast images showing cell morphology at each stage of differentiation from hiPSCs (Ff-I14s04) through mid-primitive streak and lateral plate mesoderm to ExpLBM. Scale bars: 200 µm. (B) Immunofluorescence staining of PRRX1 and SOX9 in Ff-I14s04-derived ExpLBM, with DAPI counterstaining. Scale bars: 200 µm. (C) Representative flow cytometric histograms of ExpLBM showing high expression of CD90 and CD140B.

Suppl. Fig2

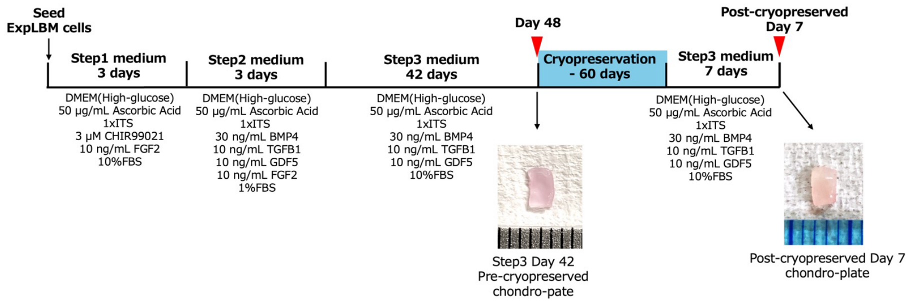


**Suppl. Fig. 2. Schematic diagram of the culture protocol for induction of chondro-plates from ExpLBM using the CAT.**

ExpLBM were seeded and sequentially cultured in Step 1, Step 2, and Step 3 media for 48 days to generate scaffold-free chondro-plates. On Day 48, the chondro-plates were cryopreserved for 60 days. After thawing, the chondro-plates were cultured in Step 3 medium for an additional 7 days before being used for transplantation.
